# Temperature and elemental sulfur shape microbial communities in two extremely acidic aquatic volcanic environments

**DOI:** 10.1101/2020.09.24.312660

**Authors:** Diego Rojas-Gätjens, Alejandro Arce-Rodríguez, Fernando Puente-Sánchez, Roberto Avendaño, Eduardo Libby, Raúl Mora-Amador, Keilor Rojas-Jimenez, Paola Fuentes-Schweizer, Dietmar H. Pieper, Max Chavarría

**Affiliations:** Centro Nacional de Innovaciones Biotecnológicas (CENIBiot), CeNAT-CONARE, 1174-1200, San José, Costa Rica; Microbial Interactions and Processes Research Group, Helmholtz Centre for Infection Research, 38124, Braunschweig, Germany; Systems Biology Program, Centro Nacional de Biotecnología (CNB-CSIC), C/Darwin 3, 28049 Madrid, Spain; Escuela de Química, Universidad de Costa Rica, 11501-2060, San José, Costa Rica; Escuela Centroamericana de Geología, Universidad de Costa Rica, 11501-2060, San José, Costa Rica; Laboratorio de Ecología Urbana, Universidad Estatal a Distancia, 11501-2060, San José, Costa Rica; Escuela de Biología, Universidad de Costa Rica, 11501-2060, San José, Costa Rica; Centro de Investigación en Electroquímica y Energía Química (CELEQ), Universidad de Costa Rica, 11501-2060, San José, Costa Rica; Centro de Investigaciones en Productos Naturales (CIPRONA), Universidad de Costa Rica, 11501-2060, San José, Costa Rica

**Keywords:** Costa Rica, Poas Volcano, Agrio River, Acidophiles, *Leptospirillum*, *Sulfobacillus*, Thermoplasmatales

## Abstract

Aquatic environments of volcanic origin provide an exceptional opportunity to study the adaptations of microbial communities to early planet life conditions such as high temperatures, high metal concentrations, and low pH. Here, we characterized the prokaryotic communities and physicochemical properties of seepage sites at the bottom of the Poas Volcano crater and the Agrio River, two geologically related extremely acidic environments located in the Central Volcanic mountain range of Costa Rica. Both locations hold a very low pH (pH 1.79-2.20) and have high sulfate and iron concentrations (Fe = 47-206 mg/L, SO_4_^2-^ = 1170-2460 mg/L measured as S), but significant differences in their temperature (90.0–95.0°C in the seepages at Poas Volcano versus 19.1–26.6 °C in Agrio River) and in the abundance of elemental sulfur. Based on the analysis of 16S rRNA gene sequences, we determined that *Sulfobacillus spp*., sulfur-oxidizing bacteria, represented more than half (58.4–78.4%) of the sequences in Poas Volcano seepage sites, while Agrio River was dominated by the iron- and sulfur-oxidizing *Leptospirillum* (7.4–55.5%) and members of the archeal order Thermoplasmatales (16.0-58.2%). Both environments share some chemical characteristics and part of their microbiota, however the temperature and the presence of reduced sulfur are likely the main distinguishing feature ultimately shaping their microbial communities. Our data suggest that in the Poas Volcano-Agrio River system there is a common metabolism but with specialization of species that adapt to the physicochemical conditions of each environment.

## Introduction

Aquatic systems of volcanic origin are ecosystems where life encounters extreme conditions and hence represent exciting habitats to understand life adaptation to harsh environments. Some volcanic lakes are extremely acidic, even reaching negative pH values as a result of the dissolution of magmatic gases and are termed hyperacid lakes [1,2], Acidity from high concentrations of sulfuric acid in hyperacid lakes arises from the high-temperature disproportionation of SO_2_ gases in the deep volcanic hydrothermal system according to reaction 3 SO_2_ +2 H_2_O → 2 H_2_SO_4_ + S [3]. The sulfuric acid is expelled into the lake where it mixes with meteoric water, whereas the produced sulfur forms a viscous molten plug at the bottom of the lake that is proposed to exert control over the eruptive activity. In Costa Rican hyperacid lakes it has been reported that magmatic degasification also contributes to acidification by introducing HCl and HF to the water [3]. Unlike sulfide-rich geothermal lakes, hydrogen sulfide is not abundant in hyperacid lakes, possibly due to reactions such as SO_2_ + 2 H_2_S → 3 S + 2 H_2_O that consume it. [3,4]

In addition to a low pH, volcanic lakes (and associated water bodies) are characterized by high concentrations of metals, such as iron, aluminum, copper, zinc, and arsenic [5–7], which are toxic to most organisms [8]. These metals derive from sulfuric acid rock dissolution or from the magma itself [1,9]. Although considerable geological and chemical characterization of these extreme environments is available [10,11], little is known about their microbial composition and ecology. The microorganisms reported inhabiting these zones are almost exclusively acidophiles, which have evolved mechanisms to survive the extreme conditions present in these ecosystems [12–15]. From a physiological perspective, acidophilic microorganisms are very diverse, comprising aerobic, facultative anaerobic, chemolithotrophic, and heterotrophic metabolism [16]; nevertheless, most of them are chemolithotrophic and oxidize reduced sulfur compounds or ferrous iron [17,18]. Despite their high metabolic versatility, these habitats usually show a very low diversity with communities mainly governed by few species [19,20].

Among the volcanic environments subjected to microbiological analysis are the Copahue–Caviahue system in Argentina, as well as the Sucio River, San Cayetano and Borbollones in Costa Rica [21–25]. Their microbial and chemical composition resembles in some aspects those found in Acid Mine Drainage (AMD) and Acid Rock Drainage (ARD) sites but are distinct in the mostly abiotic origin of the acid and soluble element composition. Recently, we have studied this difference and reassigned this type of environment as VARD (Volcanic Influenced Acid Rock Drainage) to highlight the volcanic influence [24], Culture-dependent and independent studies indicated the microbial communities present in those environments to be dominated by members of the genera *Acidithiobacillus, Leptospirillum, Thiolava* and *Gallionella* [25–28]. In addition, archaea such as Sulfolobales and Thermoplasmatales were frequently present and constituted important members of these communities, particularly in zones with high temperature [29].

The volcanic arc of Costa Rica is a mountain chain derived from the subduction of the Cocos tectonic plate under the Caribbean plate. Three of its main mountain ranges, i.e. Guanacaste, Tilaran, and Central, have active volcanoes [30]. The Poas Volcano massif, located on the Central volcanic mountain range, in Alajuela province (Figures 1A and 1B) owns one of the two hyperacid lakes found in Costa Rica. The massif reaches an altitude of 2708 m and has two crater lakes, the cold-water Lake Botos at an altitude of 2580 m and the active crater’s hot, hyperacid lake “Laguna Caliente” (Hot Lake) at about 2300 m (Figures 1C and 2). The Volcano has experienced a long period of volcanic activity alternating between states of dormancy and high activity [31,32], Multiple studies have shown its variable and dynamic geochemical behavior [33–35]. Detailed work by Rowe [31–36] on its hydrologic structure indicates that water from the Hot Lake leaks underground and resurfaces in the Agrio River (in Spanish Río Agrio) drainage basin (Figures 1D and 2). Water leaks with a 3-17 years residence time through a highly porous hydraulic conduit formed by the contact between older lava-lahar deposits and more recent active crater lava deposits. Of note, there are three sulfate chloride acidic springs on the Agrio’s drainage basin located at altitudes of 1520-2000 m on the NW flank of Poas [31].

**Figure 1.**
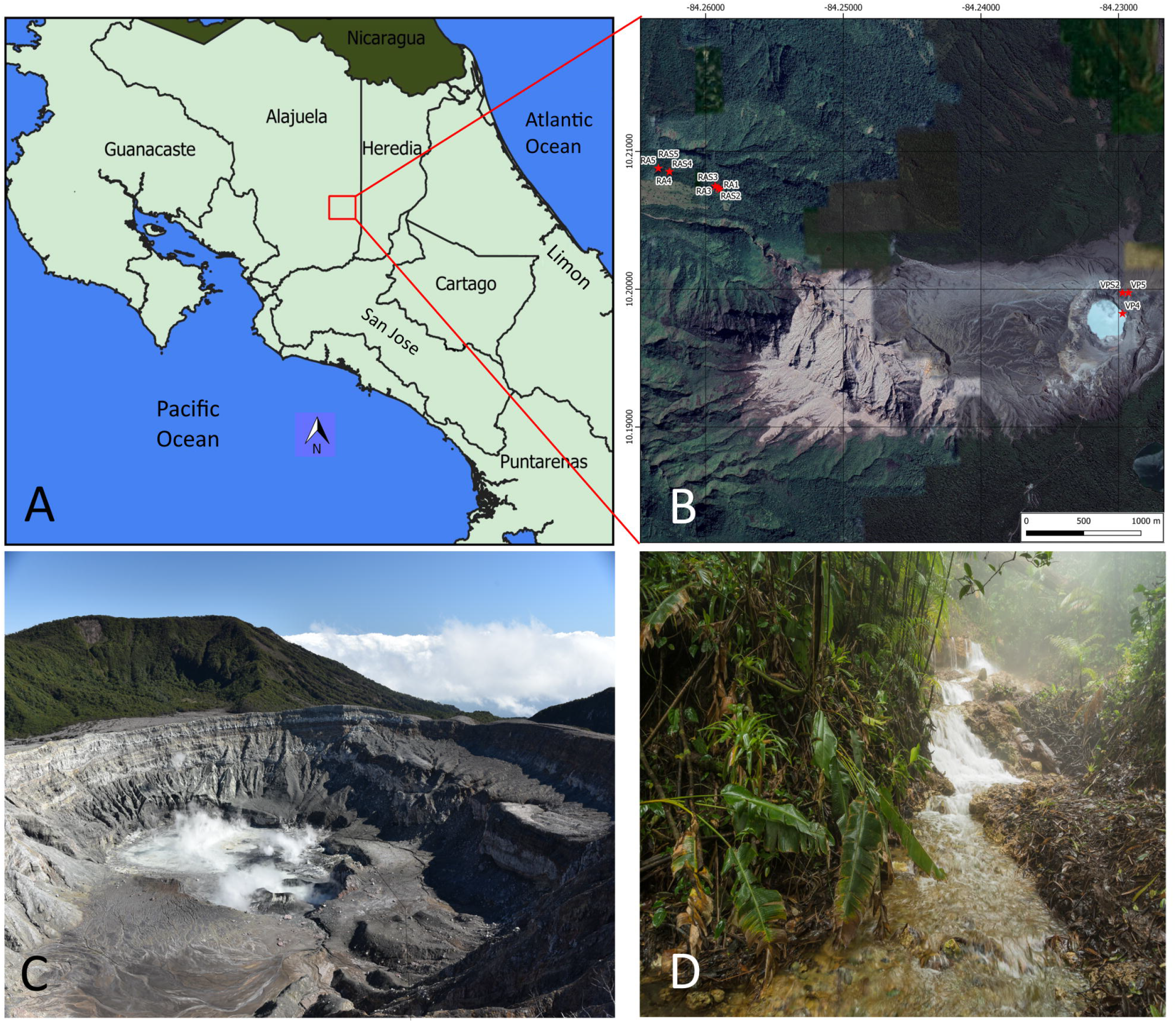
Agrio River and Poas Volcano, Alajuela, Costa Rica. A) Both places are located in the Central volcanic mountain rage, specifically in Alajuela province. B) The figure shows the Poas Volcano crater and Agrio River, the sampling points are marked with a red star. Photo taken from Google Earth Pro 7.7.3.7699.2020. Poas Volcano and surrounds 10°11’53.16” N,84°13’49.66”W, elevation 2340 M. 2D map, viewed 20 February 2020. http://www.google.com/earth/index.html. C) Poas Volcano was in a period of high instability during the sampling campaign, the figure shows a significant amount of gases and that the Lake was almost completely dry. D) Agrio River is an ARD-like habitat which shows a yellow color in sediments. The waters looked completely colorless and with little material in suspension.

**Figure 2.**
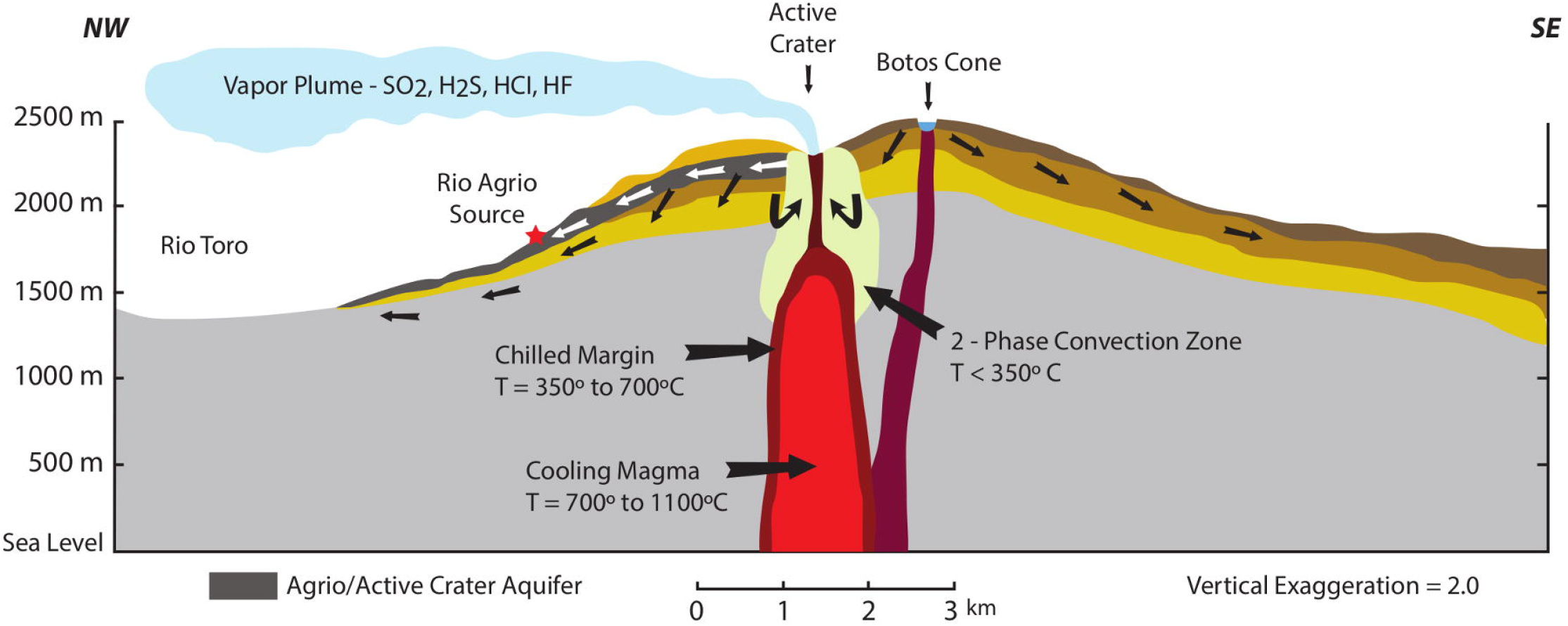
Schematic cross-section through Poas Volcano redrawn from the literature [31] illustrating the hydrologic structure of the volcano and the location of the Agrio River SO_4_-CI acidic source. The paleyellow zone underneath the active crater indicates the likely region of hydrothermal fluid convection. The darker gray layer with white arrows represents the strata associated with the dacitic andesite-lahar sequence that constitute the Agrio River aquifer and the arrows represent water transfer direction.

Far fewer studies have been conducted on the microbiota that inhabits the Poas Volcano. This is of great interest because this site is very changeable and considered one of the most extreme environments on Earth. To the best of our knowledge, only two studies have been conducted to characterize the microbial communities of the Poas Volcano and both were performed on samples taken from the hot lake in 2013 and 2017, before a strong volcanic event that took place on April 2017 (see below). These works report microbial communities dominated by species of the bacterial genus *Acidiphilium* [20] and the phylum Aquificae [37] respectively, which demonstrate the variability of the microbiota in the Poas Volcano that responds to the physicochemical conditions of this highly dynamic ecosystem. To date there are no studies carried out on the microbiota that inhabits the acidic springs near the Poas Volcano, such as the Agrio River basin.

Volcanic activity of Poas Volcano increased in early April 2017 generating frequent eruptions that eventually caused the Hot Lake to completely dry out and be replaced with a much smaller and intermittent steaming lake. The drying of the lake also exposed small seeps at the bottom of the crater (see supplementary video S1). All these dramatic changes in the physicochemical conditions at Poas Volcano and its associated water bodies should ultimately modify the microbiota. Therefore, in this study we investigated the chemical composition as well as the microbial community structure (bacteria and archaea) of the remaining extremely acidic aquatic environments of Poas Volcano. These are the hot seepage sites at the bottom of the Poas Volcano crater revealed by the drying-out of the lake (Figure S1 and supplementary video S1) and the cold source of Agrio River (Figures 1D and S2). The sites differed greatly in their temperature and the availability of elemental sulfur (S’). The sampling campaigns for this study were carried out between January and March 2018 during a brief interruption of the volcanic activity. Only a few weeks later the remains of Lake Poas completely dried out, evidencing once again the dramatic forces that shape this extreme habitat.

## Materials and Methods

### Study site

On April 12, 2017, Poas Volcano started a period of intense phreatomagmatic activity with significant eruptions that caused damage to the National Park installations and that consequently led to its closing to the public. During this period, lake Poas mostly dried out, and only during periods of rainfall a much smaller body of water was present. The sampling campaign was carried out in March 2018. The logistics for sampling in Poas Volcano were complex and extremely dangerous since, at that time, the volcano was in a period of instability. By the date of sampling, the Hot Lake of Poas was nearly dry, and it was possible to observe some areas of the crater’s bottom completely exposed (see Figures 1C, S1 and Supplementary Video S1). Upon descent to the bottom of the crater, we approached some seepage sites that flowed down to the shallow and muddy lake. In supplementary Figure S1A the red circle indicates the exact area where the samples from seepage sites were collected. Water was at or near boiling temperature at the outlet (see supplementary Figure S1D and Supplementary Video S1). However, after about 20 meters from the sampling site, the ground turned into soft and hot acidic mud, which prevented us from reaching the lake.

The source of Agrio River is on a rocky outcrop within the forest where warm underground water emerges (see Figures 1D and S2). From there, the water flows for about 7 km until its confluence with the Toro River. The river water near the source was clear and colorless, and the riverbed rocks were bare and profoundly covered with a silica-rich surface. This primary Agrio River water source has a water flow of 115 L/s and was sampled at five sites along the stream. There are two other much smaller (<5 L/s) springs. One is downstream from our sampling area at 1520 m elevation and is reported [31] as having a composition similar to that of the main source. The second one at 2000 m of altitude is a more acidic (pH 1.46) and hotter (~56°C) source described by Rowe as “El Afluente” [31,36,38]. We could not sample this latter site because, according to aerial images, it lies at the base of a cliff inside thick forest with no access trails.

### Sampling and field measurements

Poas Volcano samples were collected in March 2018. From the accessible area at the bottom of the crater (supplementary Figure S1) (the seepage sites), two composite samples of water were taken (codes VP4 and VP5). Each of these composite samples is the product of mixing three individual 1L samples. The water sampling points (VP4 and VP5) were separated by approximately 20 meters. In addition, seven sediment samples (codes VPS1, VPS2, VPS3, VPS6, VPS7, VPS8, and VPS9) at different locations in a total area of ~400 m^2^ were taken. The sediments were taken at depths not exceeding 15 cm and approximately 50 g were placed in sterile 50 mL centrifuge tubes. As explained below, only samples VP4, VP5, and VPS2 could be evaluated. The difficult access and possible danger made it impossible to take more samples of water and sediments as well as to extend the sampling to a larger area.

The sampling in the Agrio River was carried out in January 2018. Water samples were collected at five different points chosen according to their water flow characteristics along the stream (see coordinates in Table 1, Figures 1B, 1D and S2). At each location, three water samples (one liter each) were collected at different areas across the width of the river and pooled to give five composite samples. RA1 was obtained from the origin whereas sampling sites RA2, RA3, RA4, and RA5, were 45, 75, 430, and 530 m downstream, respectively, from RA1. Sediment samples were taken as reported above at each selected point (RAS2, RAS3, RAS4, and RAS5) except for location 1 (the origin) because it was a rocky site, and sediment collection was not possible.

**Table 1.** Geographic location, physical properties and chemical composition of Agrio River and Poas Volcano samples.

| Parameter | VP4 | VP5 | RA1 | RA2 | RA3 | RA4 | RA5 |
| --- | --- | --- | --- | --- | --- | --- | --- |
| Coordinates | 10°19'82.3" | 10°19'97.4" | 10°12'26.1" | 10°12'26.5" | 10°12'26.9" | 10°12'30.7" | 10°12'31.6" |
|  | 84°22'97.2" | 84°22'92.9" | 84°15'32.3" | 84°15'32.8" | 84°15'33.5" | 84°15'45.5" | 84°15'48.4" |
|  | 10.33953° | 10.34372° | 10.20725° | 10.20736° | 10.20853° | 10.20853° | 10.20878° |

|  | -84.39367° | -84.39247° | -84.25897° | -84.25911° | -84.25931° | -84.26264° | -84.26344° |
| --- | --- | --- | --- | --- | --- | --- | --- |
| Distance from Agrio River source, m | ~3300 | ~3300 | 0 | 43 | 73 | 428 | 526 |
| Altitude (M) | 2331 | 2334 | 1867 | 1859 | 1852 | 1741 | 1715 |
| Temperature (°C) / ± 0.1 | 90.0 | 95.0 | 23.6 | 23.4 | 23.3 | 19.9 | 19.1 |
| Dissolved oxygen (mg/L)/ ± 0.01 [Saturation %] | NM | NM | 4.96 [58] | 6.44 [76] | 6.88 [80] | 7.71 [84] | 7.59 [82] |
| pH ± 0.05 | 2.20 | 2.10 | 1.79 | 1.79 | 1.79 | 1.85 | 1.90 |
| Ammonium (mg/L) ± 0.03 | 0.10 | 0.10 | 0.10 | 0.10 | 0.10 | 0.10 | ND |
| Nitrate (mg/L) ± 0.05 | 0.50 | 0.40 | 0.10 | ND | 0.10 | 0.10 | ND |
| Calcium (mg/L) ± 4 | 275 | 273 | 92 | 93 | 91 | 87 | 75 |
| Magnesium (mg/L) ± 3 | 41 | 41 | 46 | 47 | 46 | 44 | 38 |
| Potassium (mg/L) ± 0.4 | 3.2 | 3.2 | 18.7 | 18.8 | 18.6 | 17.6 | 15.8 |
| Phosphorus (mg/L) ± 0.1 | 1.0 | 1.0 | 0.8 | 0.9 | 0.9 | 0.9 | 0.6 |
| Iron (mg/L) ± 4 | 206 | 205 | 63 | 63 | 63 | 59 | 47 |
| Zinc (mg/L) ± 0.05 | 0.30 | 0.30 | 0.20 | 0.20 | 0.20 | 0.20 | 0.20 |
| Copper (mg/L) | ND | ND | ND | ND | ND | ND | ND |
| Manganese (mg/L) ± 0.2 | 2.2 | 2.2 | 2.1 | 2.1 | 2.1 | 2.0 | 1.8 |
| Sodium (mg/L) ± 3 | 65 | 67 | 54 | 54 | 54 | 51 | 45 |
| Sulphur (mg/L) ± 80 | 790 | 820 | 480 | 480 | 480 | 450 | 390 |
| Electrical conductivity (mS/cm) ± 0.1 | 6.2 | 6.1 | 7.3 | 7.4 | 7.4 | 7.0 | 6.1 |

Temperature and dissolved oxygen (DO) of water samples were measured in the field with a dissolved oxygen meter YSI Model 550A (Yellow Springs Instrument Company Inc, Ohio, USA). In Poas Volcano, due to the high temperature, the use of a dissolved oxygen meter was not possible. Therefore, the temperature was measured with a partial immersion thermometer. Water samples for microbial community analysis were collected in clean and sterile bottles and processed within less than 24 h. Water samples for chemical analysis were stored at 4 °C until analysis. All the needed permits for sampling water and sediments were requested by The Institutional Commission of Biodiversity of the University of Costa Rica (resolution N° 066) and the Central Conservation Area (ACC) of the National System of Conservation Areas (SINAC) (resolution VS-040).

### Chemical Analysis

The content of calcium, iron, magnesium, manganese, potassium, sodium, phosphorus, zinc, and total sulfur was analyzed for all samples following the Standard Methods for the Examination of Water and Waste Water (SMWW) methodology described by APHA (ed 23 2017). The pH and conductivity of the water samples were measured by using a pH meter (Mettler Toledo Seven compact duo S213, Columbus, Ohio, USA) and a conductivity meter (Mettler Toledo Seven compact duo S213, Columbus, Ohio. USA) respectively. The probe was calibrated using a standard solution with a known conductivity. The analyses of metals were carried out based on APHA standards (method 2320B, Waltham, MA, USA) using Inductively Coupled Plasma (ICP-OES, Perkin Elmer Optima 8300). Spectrophotometric determination of ammonium and nitrate concentrations was carried out in triplicate (10 mL samples) following methods based on Tucker (2007), Bogren and Pruefer (2008) and Nelson (2008) respectively in a Flow Injection Analyzer (FIA, LACHAT QuickChem 8500 Series 2, Mill Rd, Milwaukee, USA). Before each batch of samples was analyzed, calibration curves were run for each analyte. All analyses were performed at CIA-UCR (Agronomical Research Center-University of Costa Rica) and CELEQ-UCR (Electrochemical and Chemical Energy Research Center).

### Total DNA isolation, construction of 16S rRNA gene libraries and Illumina sequencing

The three water samples from each sample point (1 L each) were pooled and filtered through a vacuum system under sterile conditions using a membrane filter (pore size 0.22 μm; Millipore, GV CAT No GVWP04700). To prevent rupture, another filter membrane (pore size 0.45 μm; Phenex, Nylon Part No AF0-0504) was placed below. The upper filter was collected and stored at −80 °C until processing. The DNA was extracted from aseptically cut pieces of the filter with a DNA isolation kit (PowerSoil®, MoBio, Carlsbad, CA, USA) as described by the manufacturer. Cell lysis was accomplished by two steps of bead beating (FastPrep-24, MP Biomedicals, Santa Ana, CA, USA) for 30 s at 5.5 m s^-1^. To process the sediments, a homogeneous sample of 500 mg was collected and DNA was extracted using the same protocol.

A 3-step PCR-approach was used to amplify the V5-V6 hypervariable regions of the 16S rRNA gene [39] PCR with primers 807F (5’-GGATTAGATACCCBRGTAGTC-3’) and 1050R (5’-AGYTGDCGACRRCCRTGCA-3’) [40] was used to enrich for target sequence. 1 μl of the first PCR reaction was directly used as template for the second amplification step. This PCR reaction (15 cycles) adds short overhangs to the amplicons using primers 807F_Illu (5’-<u>ACGACGCTCTTCCGATCT</u>GGATTAGATACCCBRGTAGTC-3’) and 1050R_Illu (5’-<u>GACGTGTGCTCTTCCGATCT</u>AGYTGDCGACRRCCRTGCA-3’). A third amplification step of 10 cycles (1 μl of the second PCR reaction in a total volume of 50 μl) added the two indices and Illumina adapters to amplicons. Samples that failed to give a PCR product were not further analysed. All PCR amplification steps were carried out with the PrimeSTAR HS DNA Polymerase (Takara, Otsu, Shigu, Japan) according to the manufacturer’s instructions. Obtained products were pooled in equimolar ratios and sequenced on Illumina MiSeq (2X300 bases, San Diego, USA).

### Bioinformatic and phylogenetic analysis of 16S rDNA amplicon data

Bioinformatic processing was performed as previously described [41], Raw reads were merged with the Ribosomal Database Project (RDP) assembler [42], obtaining overall 741,262 paired-end reads. Sequences were aligned within MOTHUR (gotoh algorithm using the SILVA reference database [43]) and subjected to preclustering (diffs=2) yielding so-called operational taxonomic units (OTUs) that were filtered for an average abundance of ≥0.001% and a sequence length ≥250 bp before analysis. OTUs were taxonomically classified into the SILVA v132 taxonomy [44] as reported by the SINA classification tool v1.2.11 [45]. Sequences classified as chloroplast, mitochondria or eukarya were removed for further analysis. Blastn was manually performed for the dominant OTUs found in each sample against the non-redundant and bacterial and archaeal 16S rRNA databases. A range of 44198-102016 of reads were obtained after all filtering process. Raw sequences were submitted to the sequence-read archive (SRA) of GenBank under accession number PRJNA663109.

The statistical analyses and their visualization were performed with the R statistical program [46] and Rstudio interface. Package Vegan v2.5-6 [47] and Phyloseq v1.30.0 [48] was used to calculate alpha diversity estimators (Shannon-index, Simpson, Pielou and observed richness) and Principal Coordinate Analysis (PCoA). Briefly, data was rarified to the sequencing depth of the sample with the lowest reads (44198 reads) followed by calculation of the diversities estimator. For calculating the Wunifrac distance, phylogenetic relationships were determined using the Phangorn v2.5.5 package, sequences were aligned with ClustalW and a phylogenetic tree was reconstructed based on a Neighbour Joining model. Then, data tables with the OTU abundances were normalized into relative abundances and then converted into a Wunifrac similarity matrix. For Bray-curtis similarity matrix, data was also normalized into relative abundances and then converted into the respective Matrix. PcoA was performed based on the Wunifrac distance. Observed richness, Simpson, Pielou and Shannon index were calculated and compared for significant differences among environments with Kruskal-Wallis test. Later if Kruskal-Wallis test present a significant difference a one-sided Dunn test was performed with the FSA package [49] in order to asses which environments present significant differences between them.

## Results and Discussion

### Physicochemical analysis of acidic environments

Agrio River source samples showed a pH of 1.79-1.90 and temperature ranging from 19.1 to 26.6 °C (Table 1) whereas Poas Volcano seep samples showed a slightly higher pH (2.10-2.20) but much higher temperatures (90.0-95.0°C). The temperature and pH values of the seepage sites at the bottom of the crater were both higher than those previously reported for the Hot Lake (Figure 3) [34–35,50–51] Highly acidic volcanic fluids may attack rocks resulting in a complex sequence of dissolutions and precipitations of secondary minerals. The composition of volcanic lakes thus not only reflects these processes but can also change considerably over time according to geothermal or volcanic activity and input of meteoric water. Indeed, the former Hot Lake has been closely monitored during the last decades in an attempt to use its composition as an indicator of volcanic activity. Figure 3 shows a log-log plot of the concentration of the main Rock Forming Elements (RFE) in the water along the vertical axis and their concentration in the average volcanic rock (andesite) on the horizontal axis. The resulting “Isosol” diagram also shows by means of sloping lines (Isosols) the grams of rock dissolved per liter of hydrothermal water if congruent (or complete) dissolution took place [52], Comparison of our data from the seepage sites at the bottom of the crater with previous reports shows that the water composition clearly deviates from that of the main lake, and the seepages not only are less concentrated in Mg, Na, and Fe but are low in K. The seep waters were boiling or nearly so, and their pH of 2 was less acidic than the values measured at the Hot Lake that generally are in the −0.5 to 0.5 range and only reach 1.5 during infrequent periods of low volcanic activity and high rainfall. The seepages have been rarely studied as they are only exposed when the lake either dries out completely or becomes nearly dry as it was the case here. A comparison of these analysis with data previously reported on seepages in 1995 shows similar trends [31]. Calcium was high in the 1995 sample, and the author assigned it to the dissolution of gypsum from the crater deposits and dome. The depletion of K was associated with precipitation of Alunite, KAl(SO_4_)_2_(OH)_6_, and is more likely to occur at the higher pH of the seepages.

**Figure 3.**
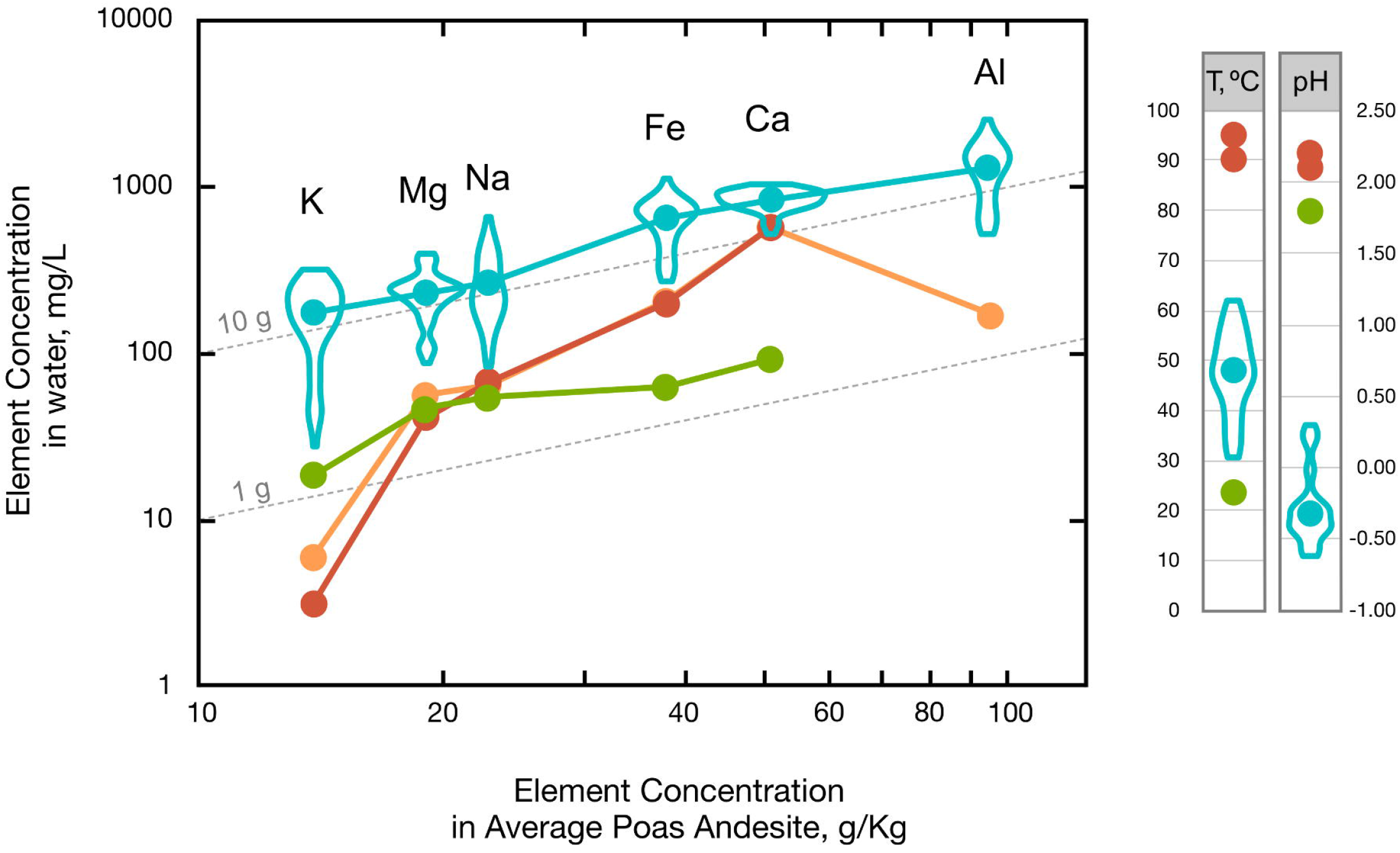
Isosol Plot of Rock Forming Element (RFE) Concentrations in water for Lake Poas, 2011-2016. Violin plots representing data distribution with average values line in blue [2], plotted against their average concentrations in Poas andesite [75]. Red dots and orange dots are data for crater seepage site VP4 (sample from this study) and from Rowe’s work respectively [31,36]. Green dots correspond to data from Agrio River Source RA1. Same color code applies for the pH and temperature data. Included are Isosol lines corresponding to dissolution of 1 g and 10 g whole rock per liter of water.

The Isosol plot also includes data for the source of Agrio River. Mass transfer calculations estimate that about 28% of the flux of RFE derives from hydrothermal inputs, and the remaining 72% comes from dissolution and leaching in the Agrio River aquifer. Concentrations seem to be affected by dilution from infiltration in the forested slopes as we sampled the accessible, more diluted, lower temperature source #25 of Rowe’s 1995 study. Despite all this, the resemblance of the Agrio River RFE profile to that of the lake suggests that the relative amounts of these elements still resemble those in the sulfate-chloride brines of the lake. There seems to be a smaller amount of Fe which could be assigned to precipitation of minerals: for example the iron mineral Jarosite is found along the Agrio River ravine [31] but water is completely clear and does not carry the large amounts of iron precipitates seen at the other VARD sites in Costa Rica studied so far [24–25], likely due to the elevated aquifer and river acidity as pH 0 water from the hyperacid crater lake only comes out at pH 2 at the Agrio River source.

Both the Agrio River and Poas water samples contain high concentrations of sulfuric acid and iron as expected in ARD, AMD or VARD systems [53–54], Even though we measured total sulfur content, considering the acidic physicochemical conditions, we assign to sulfate the sulfur that was determined in the water samples (Table 1) both in the Agrio River and Poas. As all the water samples were filtered through a 0.22 μm membrane, particulate elemental sulfur (from crater) is retained and therefore the soluble sulfur has to be in an oxidized form (i.e. sulfate). The presence of elemental sulfur was however evident throughout the sampling area as seen in the Supplementary Figures S1B, S1C and the Supplementary Video S1. The molar [iron]/[sulfur] ratios of 0.07-0.08 and 0.14-0.15 measured in the water samples are far from that expected for the oxidation of pyrite FeS2 (1:2 molar ratio Fe:S, i.e., 0.5), but consistent with what is expected for a volcanic environment, where most of the sulfate is produced by biotic and/or abiotic H_2_S and SO_2_ oxidation rather than the result of oxidation of pyrites or other metal sulfides. The formed sulfuric acid promotes the dissolution of rock minerals generating water bodies with a high content of metals (such as iron), which is consistent with our physicochemical analysis.

In summary, the physicochemical analysis of waters of Agrio River and Poas Volcano indicate that (i) both are extremely acidic environments, (ii) both contain a high content of iron and (iii) the main differentiating factors between the two environments is the higher water temperature (and thus affecting dissolved oxygen content) and availability of elemental sulfur (S^0^) in the crater.

### Analysis of microbial communities

From the nine samples collected at Poas Volcano, only three (VP4, VP5, VPS2) yielded 16S rRNA amplicons using the universal primers 807F and 1050R. The failure of samples to generate amplicons may be due to the highly extreme conditions present at Poas Volcano (pH = 2.10-2.20, Temperature = 90-95 °C, see Table 1). The extreme conditions and presence of multiple chemicals not only affect microbial diversity and biomass but also hinder the nucleic acid extraction process in the sediments and the subsequent PCR reaction. From the three samples where a successful analysis could be achieved, we identified 532 OTUs belonging to 16 phyla of domains Bacteria and Archaea (Supplementary Table S1).

The most abundant phylum in the waters of Poas Volcano seepages was Firmicutes, comprising between 55.4 and 56.3% of the sequences from each sample. Other abundant phyla included Proteobacteria (17.5-21.2%), Crenarchaeota (10.8%-11.0%), Actinobacteria (0.7-0.8%) and Nitrospirae (0.1-0.7%) (see Figure 4). The sediment sample from Poas Volcano (VPS2) was profoundly dominated by Firmicutes, representing 91.1% of the total reads. Likewise in the water samples other phyla like Proteobacteria (4.0%), Actinobacteria (1.5%), Nitrospirae (0.2%) were detected. Notably, members of the archaeal phylum Crenarchaeota were not found in the sediment sample. (See Figure 4). From all nine samples collected at the Agrio River amplicon libraries could successfully be obtained. A total of 1406 OTUs belonging to Bacteria and Archaea were identified across all samples. In the water samples, Nitrospirae (28.2-56.6%) was the dominant phylum, other abundant phyla were Euryarchaeota (16.2-20.5%), Proteobacteria (19.5-25.2%), Actinobacteria (1.6-12.7%) and unclassified bacteria or archaea (6.7-10.7%). On the other hand, Agrio River sediments samples were dominated by members of the archaeal phylum Euryarchaeota (30.4-59.1%), but Actinobacteria (10.4-21.6%), Proteobacteria (10.5-19.1%) and Nitrospirae (6.4-10.1%) were also detected (See Figure 4).

**Figure 4.**
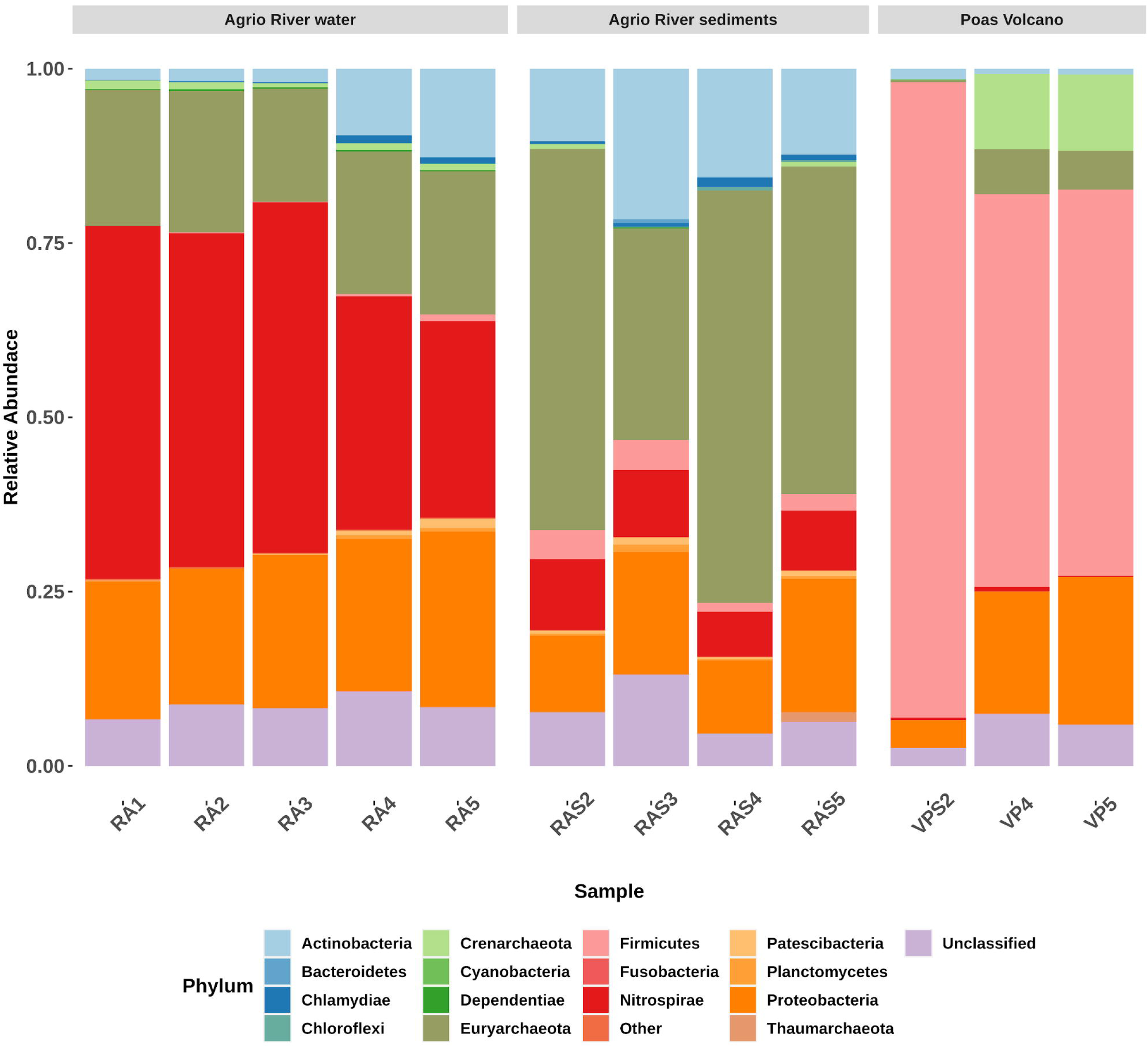
Taxonomic composition of the Poas Volcano and Agrio river microbial communities. Relative abundance of bacterial and archaeal phyla. The OTUs were taxonomically classified into the SILVA v132 taxonomy, as described in Materials and Methods. The Poas Volcano samples are termed VPS2, VP4 and VP5. Waters of Agrio River are termed RA1 to RA5 and Agrio River sediments are termed RAS2 to RAS5.

Kruskal-Wallis test showed that a significant difference between the diversity (Shannon-index and observed richness) exist between Poas Volcano, Agrio River waters and Agrio River sediments (Shannon-index p = 0.04, observed richness p =0.03). Further analysis revealed that Poas Volcano present a lower Shannon-index than both Agrio River water (One-side Dunn test, Shannon index p =0.03) and Agrio River sediments (One-side Dunn test, Shannon index p =0.03), and an observed richness lower than Agrio River water (One-side Dunn test, observed richness p =0.03) (See Supplementary Figure S3). Other indexes, such as Pielou and Simpson, don’t revealed significant differences between the environments. The lower diversity in Poas Volcano is probably associated to the high temperature conditions, which hinders life proliferation. This observation is in line with previous analysis in Poas Volcano crater [20].

In order to compare the differences between the microbial communities, Bray-Curtis similarity and Wunifrac distance were calculated. Poas Volcano samples present a low similarity according to Bray-Curtis index (18.6-19.3%), nevertheless further analysis using the Wunifrac distance, which takes into consideration the phylogenetic relationship between OTUs revealed a moderate similarity of 69.7-70.3% suggesting that sediments and water samples harbour communities with related phylotypes. Agrio River samples showed a significant difference between the community present in water and sediment samples according to Wunifrac (Similarities: 69.7-70.3, PERMANOVA p = 0.01) and Bray-Curtis distances (8.7-43.9%, PERMANOVA, p =0.008) (See Supplementary Figure S4). The difference in the microbial community between the water and sediments have been found previously in other habitats [55] and is usually produced by low oxygen concentration and nutrients diffusion in the sediments. Finally, as seen in Figure 5 a significant difference between Poas Volcano, Agrio River water and Agrio River sediments was found (PERMANOVA p = 0.002). The difference found in the microbial community structure is probably a consequence of the high temperature and sulfur concentration that is found in Poas Volcano seepages in contrast to Agrio River samples (See Table 1). As mentioned above, Poas Volcano seepages are characterized for having a high sulfur concentration and, due to its volcanic origin, most of it is in its reduced or elemental form. On the other hand, Agrio River presents less sulfur concentration, which is probably already oxidized, allowing sulfur-oxidizing bacteria to proliferate more in Poas Volcano seepages. Previous reports showed that temperature is one of the main factors that shape the structure of the microbial communities in extremely acidic habitats [56]. High temperatures in water bodies also generate a decrease in oxygen content that undoubtedly also affects the composition of the microbial community.

**Figure 5.**
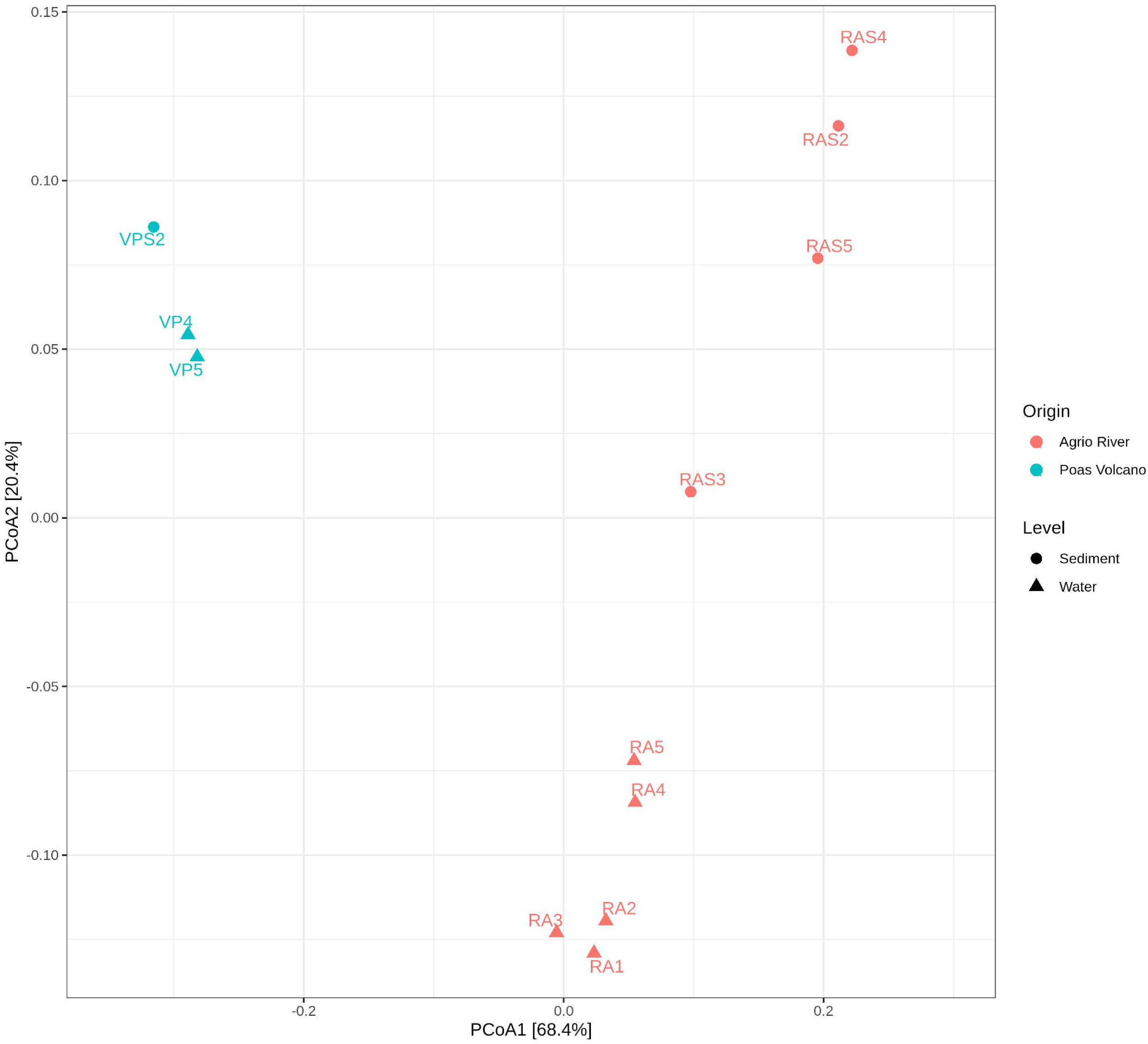
Principal Coordinate Analysis (PcoA) of the microbial communities of Poas Volcano and Agrio River. As seen, microbial communities between Poas Volcano and Agrio River differ significantly in their composition (p = 0.01). Also, it is possible to observe differences between the water column and the sediments of Agrio River (p = 0.012).

Even though significant difference in the microbial community was found, some OTUs with low abundance were shared among both places (~53% of the OTUs found in Poas Volcano were present in Agrio River samples). The presence of this OTUs in both habitats could be caused by the underground connection between the seepages of Poas Volcano and Agrio River (See Figure 2). As described above, water flow from Poas Volcano filters subsuperficially and drains in the Agrio River, dragging the microbes in the process, but due to the difference in the physicochemical conditions, the community is re-shaped in order to survive to the new niche present in the river.

The microbial communities at the seepage sites of the crater (i.e. VP4 and VP5) clearly differ from (i) previously reported for the lake [20,31] and (ii) the sediment sample obtained in this study (VPS2). Studies on samples collected in 2013 by Hynek et al. (2018) indicated Poas Volcano lake to be dominated by a unique OTU, belonging to the genus *Acidiphilium* (97% of total reads) [20] whereas samples taken in February 2017 by Fullerton et al. (2019) were dominated by bacteria from the phylum Aquificae [37], In contrast, members of the phylum Aquificae or of the genus *Acidiphilum* were absent from all three samples of seepage sites taken in March 2018. Our water samples were dominated by members of the *Sulfobacillus* genus accounting for 54-76% of sequence reads. Specifically, two bacterial OTUs were of high abundance, i.e. *Sulfobacillus* RAVP04 (15.9%) and *Sulfobacillus* RAVP06 (31.6-32.1%) (see Figure 6). They are closely related to *Sulfobacillus thermosulfidooxidans* DSM9293 (99.2%) and *Sulfobacillus acidophilus* DSM10332 (99.2%), respectively, according to the 16S rRNA sequence. Members of the *Sulfobacillus* genus have been reported to be mixotrophic, acidophilic, and thermotolerant bacteria with a versatile metabolism and accessory genes that enable them to employ multiple organic substrates as carbon sources, or to oxidize sulfur and iron and to fix CO_2_ using the Calvin pathway [57], This genus has been commonly isolated from acidic volcanic environments, containing high concentrations of reduced sulfur species [58–59]. This may indicate that the *Sulfobacillus* spp. identified in the seepages are involved mostly in sulfur and iron metabolism. Previous studies have determined that microorganisms of this genus oxidize elemental sulfur (S^0^) to produce sulfate (SO_4_^-2^) as their final product, although pathways and genetic organization to accomplish this process vary within every strain [60]. The presence of *Sulfobacillus* matches with the observed richness of elemental sulfur (supplementary Figures S1B and S1C), and the high concentrations of sulfate determined in chemical analyses (Table 1). *Sulfobacillus* spp. have mostly been reported to be present in environments with moderately high temperatures (> 65 °C) [59–62], However, it has been shown that these bacteria experience plenty of horizontal gene transfer processes, especially from archaea, which allow them to adapt to stressful conditions, such as the presence of toxic metals and shifts in temperature [63]. This condition presumably allows this genus to survive the adverse conditions present in the seepages.

**Figure 6.**
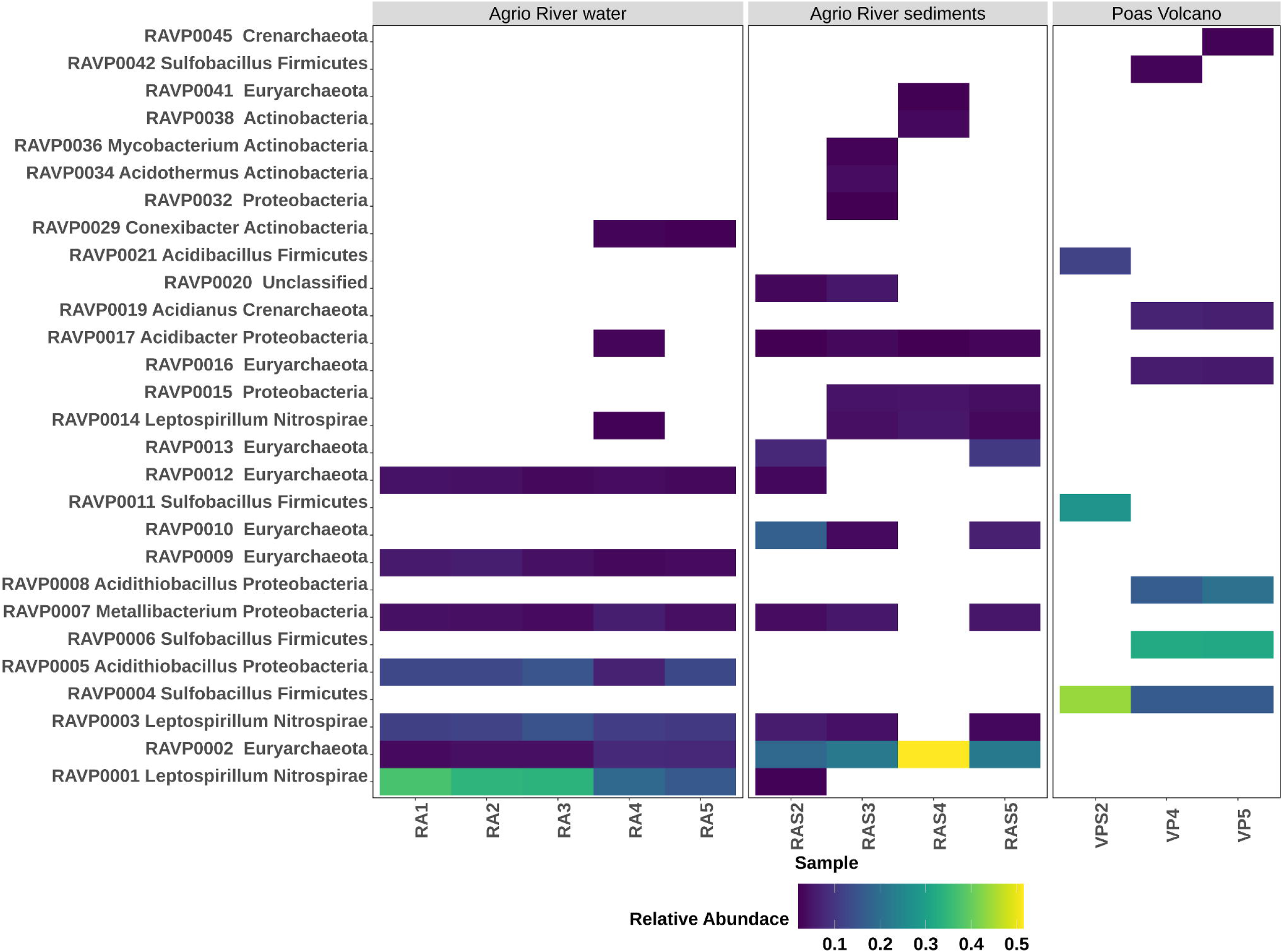
Heat map and phylogeny of the most abundant OTUs in each sample. The heat map depicts the relative percentage of 16S rRNA gene sequences assigned to each ASV (y axis) across the 11 samples analyzed (x axis).

High abundances of *Acidithiobacillus* (16.8-20.7%) were also found in the waters of Poas Volcano seepages (Figure 6). Members of the *Acidithiobacillus* genus are commonly associated with sulfur and iron metabolism, obtaining metabolic energy from the oxidation of ferrous iron, sulfide minerals (such as pyrite, chalcopyrite and metal sulfides) and inorganic sulfur compounds, to support autotrophic carbon dioxide fixation [64]. As mentioned above, the sediment of the Poas Volcano (VPS2) differ from the water samples (VP4 and VP5) (See Supplementary Figure 4). However, they share as the majority microorganism the genus *Sulfobacillus* (RAVP04 43.9%; RAVP11 27.5%) as one of the highly abundant microorganisms. A member of the genus *Acidibacillus*(RAVP21, 11.9%), which was not found in the Poas Volcano water samples (i.e. VP4 and VP5) was found as the second most abundant genera in the sediment samples (VPS2). To date, the genus *Acidibacillus* reports two species: *Acidibacillus ferrooxidans* and *Acidibacillus sulfuroxidans*, which are classified as iron and sulfur-oxidizing bacteria, isolated from geothermal sites [65]

In Agrio River, *Leptospirillum* was found as the dominant microorganism the water column (28.2-50.6%), and was also highly abundant in the sediments (6.4-10.1%). Members of the *Leptospirillum* genus are known as aerobic chemolithoautotrophic and acidophilic iron oxidizers [66]. They resemble fastidious bacteria, which are only able to grow using iron minerals (such as pyrite or other Fe^+2^ sources) and in some cocultures grow with some sulfur related species [67]. The high quantity of *Leptospirillum* in Agrio River samples is probably due to its elevated iron concentration, which, altogether with the low pH, produce an appropriate niche for members of this genus.

On the other hand, members of Thermoplasmatales dominated in the sediment samples (30.0-58.2%) and were somewhat less abundant in the water column (16.0-20.3%) (see Figure 6). *Acidithiobacillus* was also of high abundance in the water column (16.3-20.18%) but poorly detected in sediment (<1.4%) whereas Acidimicrobia and Acidithiobacillaceae RCP1-48 were more abundant in sediments (7-10% and 2.6-6.2%) compared to the water column (1.0-6.1% and 0.8-2.0%). Interestingly, the *Sulfobacillus* that was found as the dominant OTU in seepage samples from Poas Volcano was found in low abundance (<2.2% in sediments and <0.05% in water column). Thermoplasmatales RAVP02 was the dominant OTU (18.1-50.4%) particularly in sediments samples. However, its phylogeny could not be further determined. Members of the Thermoplasmatales order usually are isolated from sulfuric environments with low pH [23,68–69] and have been reported to be involved in sulfur oxidation and reduction [70–71].

Taking together the chemical and microbiological data, we suggest that the difference in the prokaryotic communities of both places can be mainly associated with differences in (i) temperature which also affects the dissolved oxygen content and (ii) the presence of abundant elemental sulfur in Poas Volcano. Similar effects had been reported in other extremely acidic environments [56–72], where temperature is one of the significant properties that shift the microbial composition. It is also important to consider that the seepages are in a volcanic crater environment with only inorganic electron donors in contrast to the Agrio River aquifer and stream which lie inside a forest where both infiltration to the aquifer and runoff into the stream inoculate organic matter and likely contribute to the difference in the community of both places. Additionally, differences were observed between the water and sediment samples both in the Agrio River and in the Poas Volcano, the difference in the microbial community is probably associated to the oxygen and nutrient availability as has been shown in other aquatic environments.

## Concluding Remarks

The characterization of the microbial communities from Poas Volcano and Agrio River unravel part of the acidophilic microbial diversity in Costa Rica’s volcanic influenced ecosystems. The two studied environments are geologically related, since it has been previously shown that the waters of Poas Volcano move underground and emerge relatively fast, in a few years, in the Agrio River basin. As a consequence, both water sources show comparable physicochemical properties typical for volcanic habitats, like a very low pH and high sulfate and iron concentrations, but differ sharply in temperature and the presence of elemental sulfur. These two related sources of acidic volcanic water fulfill our proposed definition of VARD [24], At the volcanic environment of the crater we found seepages with water at high temperatures (and therefore low oxygen levels). In the Agrio River basin the same source of water from the crater becomes now cold and oxic. Both environments have chemical similarities but significant biological differences even though they are subsuperficially connected. The difference is caused by two factors that shape and giving specificity to the microbial community of each ecosystem; these are the temperature and the presence of reduced sulfur (mainly S^0^). The high temperature and availability of sulfur at the crater likely favors the growth of thermoacidophilic and sulfur- and iron-oxidizing bacteria such as *Sulfobacillus* or *Acidithiobacillus*. On the other hand, the lower temperature and chemical composition (i.e. high concentrations of iron, sulfate and oxygen) of the Agrio River favors the growth of mesophilic microorganisms capable of oxidizing iron and sulfur such as *Leptospirillum*. Our data suggest that both ecosystems present a similar biochemistry and metabolism based on the oxidation of iron and sulfur, however, temperature and the presence of elemental sulfur shape the actors that participate in these processes. Our data gives insights into understanding the diversity and taxonomic composition of microbial communities in acidic environments from Costa Rica and fits the idea that extreme habitats present species specialization but a relatively general metabolic pattern across samples. Similar results are found in the microbiome of other niches [73–74], where taxonomic data is very variable, but metabolic pathways present have a relatively constant composition across different samples and locations.

## Acknowledgements

We thank Sergio Paz and Gerardo Chavarría for contributing and operating the drones in the sampling campaign at the Poas Volcano. We also thank Solange Voysest for her help in figure preparation.

## Author contributions

AAR, FPS, EL, MC conceived and designed the experiments; AAR, RA, RMA, DRG, PF performed the experiments; AAR, FPS, DRG, KR, MC analyzed the data; PF, MC, DHP contributed reagents or materials or analytical tools; DRG, PF, EL, KR, DHP, MC wrote the paper. All authors reviewed and approved the final version of the manuscript.

## References

1. Rouwet D, Mora-Amador R, Ramírez-Umaña CJ, González G, Inguaggiato S (2016) Dynamic fluid recycling at Laguna Caliente (Poás, Costa Rica) before and during the 2006–ongoing phreatic eruption cycle (2005— 10). Geological Society 437:73.–96. https://doi.org/10.1144/SP437.11

2. Rouwet D, Mora-Amador RA, Sandri L, Ramírez-Umaña G, González G, Pecoraino G, Capaccioni B (2019) 39 years of geochemical monitoring of Laguna Caliente crater lake, Poas: Patterns from the past as keys for the future. In: Tassi F., Vaselli O., Mora Amador R. (eds) Poás Volcano. Active volcanoes of the World. Springer, Cham

3. Delmelle P, Bernard A (2015) The remarkable chemistry of sulfur in hyper-acid crater lakes: A scientific tribute to Bokuichiro Takano and Minoru Kusakabe. In: Rouwet D, Christenson B, Tassi F, Vandemeulebrouck J (eds) Volcanic Lakes. Advances in Volcanology. Springer, Berlin, Heidelberg

4. Mora Amador RA, Rouwet D, Vargas P, Oppenheimer C (2019) The extraordinary sulfur volcanism of Poas from 1828 to 2018. In: Tassi F, Vaselli O, Mora Amador R (eds) Poás Volcano. Active Volcanoes of the World. Springer, Cham. Switzerland

5. Dongarrà G, Varrica D (1998) The presence of heavy metals in air particulate at Vulcano island. Sci. Total Environ 212(1):1–9.https://doi.org/10.1016/S0048-9697(97)00323-9

6. Vigneri R, Malandrino P, Gianì F, Russo M, Vigneri P (2017) Heavy metals in the volcanic environment and thyroid cancer. Mol Cell Endocrinol 457:73–80. https://doi.Org/10.1016/j.mce.2016.10.027

7. Dœlsch E, Macary HS, Van de Kerchove V (2006) Sources of very high heavy metal content in soils of volcanic island (La Réunion). J. Geochem. Explor 88:1–3. https://doi.org/10.1016/j.gexplo.2005.08.037

8. Tchounwou PB, Yedjou CG, Patlolla AK, Sutton DJ (2012) Heavy metal toxicity and the environment. In: Luch A (eds) Molecular, clinical and environmental toxicology. Springer, Basel

9. Bradl HB (2005) Sources and origins of heavy metals. In: Bradl HB (eds) Heavy metals in the environment: origin, interaction and remediation, Elsevier

10. Sakai H, Casadevall TJ, Moore JG (1982) Chemistry and isotope ratios of sulfur in basalts and volcanic gases at Kilauea volcano, Hawaii. Geochimica et Cosmochimica Acta 46(5):729–738. https://doi.org/10.1016/0016-7037(82)90024-2

11. Malahoff A, McMurtry G, Wiltshire J, Yeh HG (1982) Geology and chemistry of hydrothermal deposits from active submarine volcano Loihi, Hawaii. Nature 298:234–239. https://doi.org/10.1038/298234a0

12. Norris PR (2007) Acidophile diversity in mineral sulfide oxidation. In: Rawlings DE, Johnson DB (eds) Biomining. Springer, Berlin, Heidelberg, https://doi.org/10.1007/978-3-540-34911-2_10

13. Dopson M, Holmes DS (2014) Metal resistance in acidophilic microorganisms and its significance for biotechnologies. Appl Microbiol Biotechnol 98:8133–814. https://doi.org/10.1007/s00253-014-5982-2

14. Reed CJ, Lewis H, Trejo E, Winston V, Evilia C (2013) Protein adaptations in archaeal extremophiles. Archaea. 2013:e373275. https://doi.org/10.1155/2013/373275

15. Dopson M, Johnson DB (2012) Biodiversity, metabolism and applications of acidophilic sulfur-metabolizing microorganisms. Environ Microbiol. 14(10):2620–2631. https://doi.org/10.1111/j.1462-2920.2012.02749.x

16. Mirete S, Morgante V, González-Pastor JE (2017) Acidophiles: diversity and mechanisms of adaptation to acidic environments. In: Stan-Lotter H, Fendrihan S (eds) Adaption of microbial life to environmental extremes. Springer, Cham.

17. Wang R, Lin JQ, Liu XQ et al (2019) Sulfur oxidation in the acidophilic autotrophic Acidithiobacillus spp. Front. Microbiol 3:e290. https://doi.org/10.3389/fmicb.2018.03290

18. Jones DS, Kohl C, Grettenberger C, Larson LN, Burgos WD, Macalady JL (2014) Geochemical niches of Fe-oxidizing acidophiles in an acidic coalmine drainage. Appl. Environ. Microbiol. 14:e02919. https://doi.org/10.1128/AEM.02919-14

19. Gómez F, Cavalazzi B, Rodríguez N et al (2019) Ultra-small microorganisms in the polyextreme conditions of the Dallol volcano, Northern Afar, Ethiopia. Sci Rep 9:e7907. https://doi.org/10.1038/s41598-019-44440-8

20. Hynek BM, Rogers KL, Antunovich M, Avard G, Alvarado GE (2018) Lack of microbial diversity in an extreme Mars analog setting: Poás Volcano, Costa Rica. Astrobiology 18(7):923–933. https://doi.org/10.1089/ast.2017.1719

21. Chiacchiarini P, Lavalle L, Giaveno A, Donati E (2010) First assessment of acidophilic microorganisms from geothermal Copahue–Caviahue system. Hydrometallurgy 104(3-4):334–341. https://doi.org/10.1016/j.hydromet.2010.02.020

22. López-Archilla A, Marin I, Amils R (2001) Microbial community composition and ecology of an acidic aquatic environment: The Tinto River, Spain. Microb Ecol 41:20–35. https://doi.org/10.1007/s002480000044

23. Arce-Rodríguez A, Puente-Sánchez F, Avendaño R et al (2019) Thermoplasmatales and sulfur-oxidizing bacteria dominate the microbial community at the surface water of a CO2-rich hydrothermal spring located in Tenorio Volcano National Park, Costa Rica. Extremophiles 23:177–187. https://doi.org/10.1007/s00792-018-01072-6

24. Arce-Rodríguez A, Puente-Sánchez F, Avendaño R et al (2020) Microbial community structure along a horizontal oxygen gradient in a Costa Rican volcanic influenced acid rock drainage system. Microb Ecol. https://doi.org/10.1007/s00248-020-01530-9

25. Arce-Rodríguez A, Puente-Sánchez F, Avendaño R et al (2017) Pristine but metal-rich Río Sucio (Dirty River) is dominated by Gallionella and other iron-sulfur oxidizing microbes. Extremophiles 21:235–243. https://doi.org/10.1007/s00792-016-0898-7

26. Fujimura R, Sato Y, Nishizawa T, Nanba K, Oshima K, Hattori M, Kamijo T, Ohta H (2012) Analysis of early bacterial communities on volcanic deposits on the island of Miyake (Miyake-jima), Japan: a 6-year study at a fixed site. Microbes Environ. 27(1):19–29. https://doi.org/10.1264/jsme2.me11207

27. Yan L, Zhang S, Yu G, Ni Y, Wang W, Hu H, Chen P (2015) Draft genome of iron-oxidizing bacterium Leptospirillum sp. YQP-1 isolated from a volcanic lake in the Wudalianchi volcano, China. Genom Data 6:164–165. https://doi.org/10.1016/j.gdata.2015.09.002

28. Danovaro R, Canals M, Tangherlini M et al (2017) A submarine volcanic eruption leads to a novel microbial habitat. Nat Ecol Evol 1:e0144. https://doi.org/10.1038/s41559-017-0144

29. Johnson DB (1998) Biodiversity and ecology of acidophilic microorganisms. FEMS Microbiol Ecol 27(4):307–317. https://doi.org/10.1111/j.1574-6941.1998.tb00547.x

30. Castellón E, Martínez M, Madrigal-Carballo S, Arias ML, Vargas WE, Chavarría M (2013) Scattering of light by colloidal aluminosilicate particles produces the unusual sky-blue color of Río Celeste (Tenorio volcano complex, Costa Rica). PLoS One 8:e75165

31. Rowe GL, Brantley SL, Fernandez JF, Borgia A (1995) The chemical and hydrologic structure of Poas Volcano, Costa Rica. J Volcanol Geotherm Res 64:233–267. https://doi.org/10.1016/0377-0273(92)90003-V

32. Ruiz P, Mana S, Gutiérrez G, Alarcón G, Garro J, Soto GJ (2018) Geomorphological insights on human-volcano interactions and use of volcanic materials in pre-hispanic cultures of Costa Rica through the holocene. Front Earth Sci 6:e13.https://doi.org/10.3389/feart.2018.00013

33. Taran Y, Tassi F, Varekamp J, Inguaggiato S, Kalacheva E (2017) Volcano-hydrothermal systems. Elsevier.

34. Rymer H, Cassidy J, Locke CA, Barboza MV, Barquero J, Brenes J, Van der Laat R (2000) Geophysical studies of the recent 15-year eruptive cycle at Poas Volcano, Costa Rica. J Volcanol Geotherm Res 97(1-4):425–442. https://doi.org/10.1016/S0377-0273(99)00166-3

35. Tassi F, Vaselli O, Mora Amador RA (2019) Poás Volcano: The pulsing heart of Central America volcanic zone. Springer International Publishing. Cham

36. Rowe GL, Brantley SL, Fernandez M, Fernandez JF, Borgia A, Barquero J (1992) Fluid-volcano interaction in an active stratovolcano: The crater lake system of Poas Volcano, Costa Rica. J Volcanol Geotherm Res 49:23–51.

37. Fullerton KM, Schreck M, Yucel M et al (2020) Tectonic processes shape biosphere-geosphere feedbacks across a convergent margin. EarhArXiv preprints, https://eartharxiv.org/gyr7n/.

38. Rowe GL, Ohsawa S, Takano B, Brantley SL, Fernandez JF, Barquero J (1992) Using crater Llake chemistry to predict volcanic activity at Poas Volcano, Costa Rica. Bull Volcanol 54(6):494–503. https://doi.org/10.1007/BF00301395.

39. Rath S, Heidrich B, Pieper DH, Vital M (2017) Uncovering the trimethylamine-producing bacteria of the human gut microbiota. Microbiome 5:e54. https://doi.org/10.1186/s40168-017-0271-9

40. Bohorquez LC, Delgado-Serrano L, López G, et al (2012) In-depth characterization via complementing culture-independent approaches of the microbial community in an acidic hot spring of the Colombian Andes. Microb Ecol 63:103–11

41. Schulz C, Schütte K, Koch N et al (2018) The active bacterial assemblages of the upper GI tract in individuals with and without Helicobacter infection. Gut 67:216–225. https://doi.org/10.1186/s40168-017-0271-9

42. Cole JR, Wang Q, Fish JA et al (2014) Ribosomal database project: data and tools for high throughput rRNA analysis. Nucleic Acids Res 42:633–642. https://doi.org/10.1093/nar/gkt1244

43. Schloss PD, Westcott SL, Ryabin T et al (2009) Introducing mothur: open-source, platform-independent, community-supported software for describing and comparing microbial communities. Appl Environ Microbiol 75:7537–7541. https://doi.org/10.1128/AEM.01541-09

44. Yilmaz P, Parfrey LW, Yarza P et al (2014) The SILVA and “All-Species Living Tree Project (LTP)” taxonomic frameworks. Nucleic Acids Res. 42:643–648. https://doi.org/10.1093/nar/gkt1209

45. Pruesse E, Peplies J, Glöckner FO (2012) SINA: accurate high-throughput multiple sequence alignment of ribosomal RNA genes. Bioinformatics 28:1823–1829. https://doi.org/10.1093/bioinformatics/bts252

46. Core Team R (2017) R: A language and environment for statistical computing. R foundation for statistical computing, Vienna, Austria http://www.R-project.org/

47. Oksanen J, Blanchet FG, Friendly M, et al (2017) Vegan: Community Ecology Package. R package Version 2.4–3. https://CRAN.R.project.org/package=vegan

48. McMurdie PJ, Holmes S (2013) phyloseq: An R package for reproducible interactive analysis and graphics of microbiome census data. PLoS One 8(4):e61217. https://doi.org/10.1371/journal.pone.0061217

49. Ogle DH, Wheeler P, Dinno A (2020). FSA: Fisheries Stock Analysis. R package version 0.8.30. https://github.com/droglenc/FSA.

50. Rodríguez A, J. van Bergen M (2017) Superficial alteration mineralogy in active volcanic systems: An example of Poas Volcano, Costa Rica. J Volcanol Geotherm Res 346:54–80.https://doi.org/10.1016/j.jvolgeores.2017.04.006

51. Martínez M, Fernández E, Valdés J et al (2000) Chemical evolution and volcanic activity of the active crater lake of Poas volcano, Costa Rica, 1993–1997. J Volcanol Geotherm Res 97:127–141. https://doi.org/10.1016/S0377-0273(99)00165-1

52. Varekamp JC (2015) The Chemical composition and evolution of volcanic lakes. In: Rouwet D., Christenson B, Tassi F, Vandemeulebrouck J (eds) Volcanic lakes. Advances in volcanology. Springer, Berlin, Heidelberg.

53. Torres-Ceron DA, Acosta-Medina CD, Restrepo-Parra E (2019) Geothermal and mineralogic analysis of hot springs in the Puracé-La Mina sector in Cauca, Colombia. Geofluids 2019:e3191454

54. Amils R, Fernández-Remolar D, The IPBSL Team (2014) Río Tinto: A geochemical and mineralogical terrestrial analogue of Mars. Life 4:511–534. https://doi.org/10.3390/life4030511

55. Sommers P, Darcy JL, Porazinska DL et al (2019) Comparison of microbial communities in the sediments and water columns of frozen cryoconite holes in the McMurdo dry valleys, Antarctica. Front Microbiol 10:e65. https://doi.org/10.3389/fmicb.2019.00065

56. Quatrini R, Johnson DB (2018) Microbiomes in extremely acidic environments: functionalities and interactions that allow survival and growth of prokaryotes at low pH. Curr Opin Microbiol 43:139–147.https://doi.org/10.1016/j.mib.2018.01.011

57. Guo X, Yin H, Liang Y et al. (2014) Comparative genome analysis reveals metabolic versatility and environmental adaptations of Sulfobacillus thermosulfidooxidans Strain ST. PLoS One 9(6): e99417. https://doi.org/10.1371/journal.pone.0099417

58. Watling HR, Perrot FA, Shiers DW (2008) Comparison of selected characteristics of Sulfobacillus species and review of their occurrence in acidic and bioleaching environments. Hydrometallurgy 93:57–65. https://doi.org/10.1016/j.hydromet.2008.03.001

59. Melamud VS, Pivovarova TA, Tourova TP et al (2003) Sulfobacillus sibiricus sp. nov., a New moderately thermophilic bacterium. Microbiology 72:605–612. https://doi.org/10.1023/A:1026007620113

60. Justice NB, Norman A, Brown CT, Singh A, Thomas BC, Banfield JF (2014) Comparison of environmental and isolate Sulfobacillus genomes reveals diverse carbon, sulfur, nitrogen, and hydrogen metabolisms. BMC Genomics 15:e1107. https://doi.org/10.1186/1471-2164-15-1107

61. Zeng W, Wu C, Zhang R, Hu P, Qiu G, Gu G, Zhou H (2009) Isolation and identification of moderately thermophilic acidophilic iron-oxidizing bacterium and its bioleaching characterization. Trans Nonferrous Met Soc China 19:222–227. https://doi.org/10.1016/S1003-6326(08)60256-3

62. Johnson DB, Joulian C, d’Hugues P, Hallberg KB (2008) Sulfobacillus benefaciens sp. nov., an acidophilic facultative anaerobic Firmicute isolated from mineral bioleaching operations. Extremophiles 12:e789. https://doi.org/10.1007/s00792-008-0184-4

63. Zhang X, Liu X, Liang Y et al (2017) Adaptive Evolution of Extreme Acidophile Sulfobacillus thermosulfidooxidans potentially driven by horizontal gene transfer and gene loss. Appl Environ Microbiol. 83(7):e03098–16. https://doi.org/10.1128/AEM.03098-16

64. Valdés J, Pedroso I, Quatrini R, Dodson RJ, Tettelin H, Blake R, Eisen JA, Holmes DS (2008) Acidithiobacillus ferrooxidans metabolism: from genome sequence to industrial applications. BMC Genomics 9:e597. https://doi.org/10.1186/1471-2164-9-597

65. Schopf S, Ullrich SR, Heine T, Schlömann M (2017) Draft genome of the heterotrophic iron-oxidizing bacterium “Acidibacillus ferroxidans” Huett2, isolated from a mine drainage ditch in Freiberg, Germany. Genome announc 5(19):e00323–17. https://doi.org/10.1128/genomeA.00323-17

66. Daims H (2014) The Family Nitrospiraceae. In: Rosenberg E, DeLong EF, Lory S et al (eds) The Prokaryotes. Springer, Berlin, Heidelberg https://doi.org/10.1007/978-3-642-38954-2_126

67. Smith SL, Johnson DB (2018) Growth of Leptospirillum ferriphilum in sulfur medium in co-culture with Acidithiobacillus caldus. Extremophiles 22:327–333. https://doi.org/10.1007/s00792-018-1001-3

68. Segerer A, Langworthy TA, Stetter KO (1998) Thermoplasma acidophilum and Thermoplasma volcanium sp. nov. from Solfatara Fields. Syst Appl Microbiol 10(2):161–171.https://doi.org/10.1016/S0723-2020(88)80031-6

69. Serour E, Antranikian G (2002) Novel thermoactive glucoamylases from the thermoacidophilic Archaea Thermoplasma acidophilum, Picrophilus torridus and Picrophilus oshimae. Antonie Van Leeuwenhoek 81:73–83. https://doi.org/10.1023/A:1020525525490

70. Golyshina OV, Lünsdorf H, Kublanov IV, Goldenstein NI, Hinrichs KU, Golyshin PN (2016) The novel extremely acidophilic, cell-wall-deficient archaeon Cuniculiplasma divulgatum gen. nov., sp. nov. represents a new family, Cuniculiplasmataceae fam. nov., of the order Thermoplasmatales. Int J Syst Evol Microbiol. 66(1):332–340. https://doi.org/10.1099/ijsem.0.000725

71. Reysenbach AL, Brileya K (2014) The family Thermoplasmataceae. In: Rosenberg E, DeLong EF, Lory S, Stackebrandt E, Thompson F (eds) The Prokaryotes. Springer, Berlin, Heidelberg.

72. Ward L, Taylor MW, Power JF, Scott BJ, McDonald IR, Stott MB (2017) Microbial community dynamics in Inferno Crater Lake, a thermally fluctuating geothermal spring. ISME J. 11(5):1158–1167. https://doi.org/10.1038/ismej.2016.193

73. Eng A, Borenstein E (2018). Taxa-function robustness in microbial communities. Microbiome 6(1):e45. https://doi.org/10.1186/s40168-018-0425-4

74. Louca S, Jacques SMS, Pires APF, Leal JS, Srivastava DS, Parfrey LW, Farjalla VF, Doebeli M (2016) High taxonomic variability despite stable functional structure across microbial communities. Nat Ecol Evol. 1(1):e15. https://doi.org/10.1038/s41559-016-0015. PMID: 28812567.

75. Cigolini C, Kudo AM, Brookins DG, Ward D (1991) The petrology of Poas Volcano Lavas: Basalt-andesite relationship and their petrogenesis within the magmatic arc of Costa Rica. J Volcanol and Geotherm Res 48(3): 367–384. https://doi.org/10.1016/0377-0273(91)90052-2.

