## Supplementary information for "Temperature and elemental sulfur shape microbial communities in two extremely acidic aquatic volcanic environments"

Diego Rojas-Gätjens^1,^ Alejandro Arce-Rodríguez^2α^, Fernando Puente-Sánchez^3^, Roberto Avendaño^1^, Eduardo Libby^4^, Raúl Mora-Amador^5,6^, Keilor Rojas-Jimenez^7^, Paola Fuentes-Schweizer^4,8^, Dietmar H. Pieper^2^ & Max Chavarría^1,4,9*^

^1^Centro Nacional de Innovaciones Biotecnológicas (CENIBiot), CeNAT-CONARE, 1174-1200, San José, Costa Rica ^2^Microbial Interactions and Processes Research Group, Helmholtz Centre for Infection Research, 38124, Braunschweig, Germany ^3^Systems Biology Program, Centro Nacional de Biotecnología (CNB-CSIC), C/Darwin 3, 28049 Madrid, Spain ^4^Escuela de Química, Universidad de Costa Rica, 11501-2060, San José, Costa Rica ^5^Escuela Centroamericana de Geología, Universidad de Costa Rica, 11501-2060, San José, Costa Rica ^6^Laboratorio de Ecología Urbana, Universidad Estatal a Distancia, 11501-2060, San José, Costa Rica ^7^Escuela de Biología, Universidad de Costa Rica, 11501-2060, San José, Costa Rica, ^8^Centro de Investigación en Electroquímica y Energía Química (CELEQ), Universidad de Costa Rica, 11501-2060, San José, Costa Rica ^9^Centro de Investigaciones en Productos Naturales (CIPRONA), Universidad de Costa Rica, 11501-2060, San José, Costa Rica.

Keywords: Costa Rica, Poas Volcano, Rio Agrio, Acidophiles, *Leptospirillum, Sulfobacillus*, Thermoplasmatales

^α^ Current affiliation:

Department of Molecular Bacteriology

Helmholtz Centre for Infection Research

38124, Braunschweig, Germany

* Correspondence to: Max Chavarría

Escuela de Química & Centro de Investigaciones en Productos Naturales (CIPRONA)

Universidad de Costa Rica

Sede Central, San Pedro de Montes de Oca

San José, 11501-2060, Costa Rica

Phone (+506) 2511 8520.  Fax (+506) 2253 5020

ORCID: <https://orcid.org/0000-0001-5901-3576>

**LEGENDS OF SUPPLEMENTARY TABLE**

**Table S1. DNA sequence and phylogenetic assignment of the most abundant phylotypes detected in Poas Volcano and Agrio River using Illumina-based amplicon deep-sequencing.**

See Excel file.

**LEGENDS OF SUPPLEMENTARY FIGURES**

**Figure S1. Poas Volcano, Alajuela, Costa Rica.** A). Exact location of the seepage sample in Poas Volcano crater B) The samples were taken during a period of high instability, with a constant expelled of gases and high temperatures. C) Yellow sulfur deposits were abundant in Poas Volcano crater. D) High temperatures were common in the seepages sites in the crater as almost all water present were boiling.

**Figure S2. Agrio River, Alajuela, Costa Rica.** A). A rocky surface is observed where Agrio River is born B) Agrio River sediments present a yellowish color.

**Figure S3. Heat map representing the similarity matrix of Agrio River and Poas Volcano samples based on Bray-Curtis and wUniFrac distances.** Distances were calculated as described in Materials and Methods

**Figure S4. Diversity measures of the samples in Agrio River and Poas Volcano**. The diversity measures (Shannon, Simpson and Observed Richness) were calculated using phyloseq. Figure shows A) diversity measures of all samples grouped by sample point. B) diversity measures of all samples.

**LEGEND OF SUPPLEMENTARY VIDEO**

**Video S1. Conditions of the Poas Volcano during March 2018**. The video shows the conditions of the Poas Volcano during the sampling campaign. The hot lagoon was in the process of drying out exposing part of the crater. The aerial photographs were taken using drones.
