## Supplementary figures and images for "Temperature and elemental sulfur shape microbial communities in two extremely acidic aquatic volcanic environments"

### Supp. Figure S1

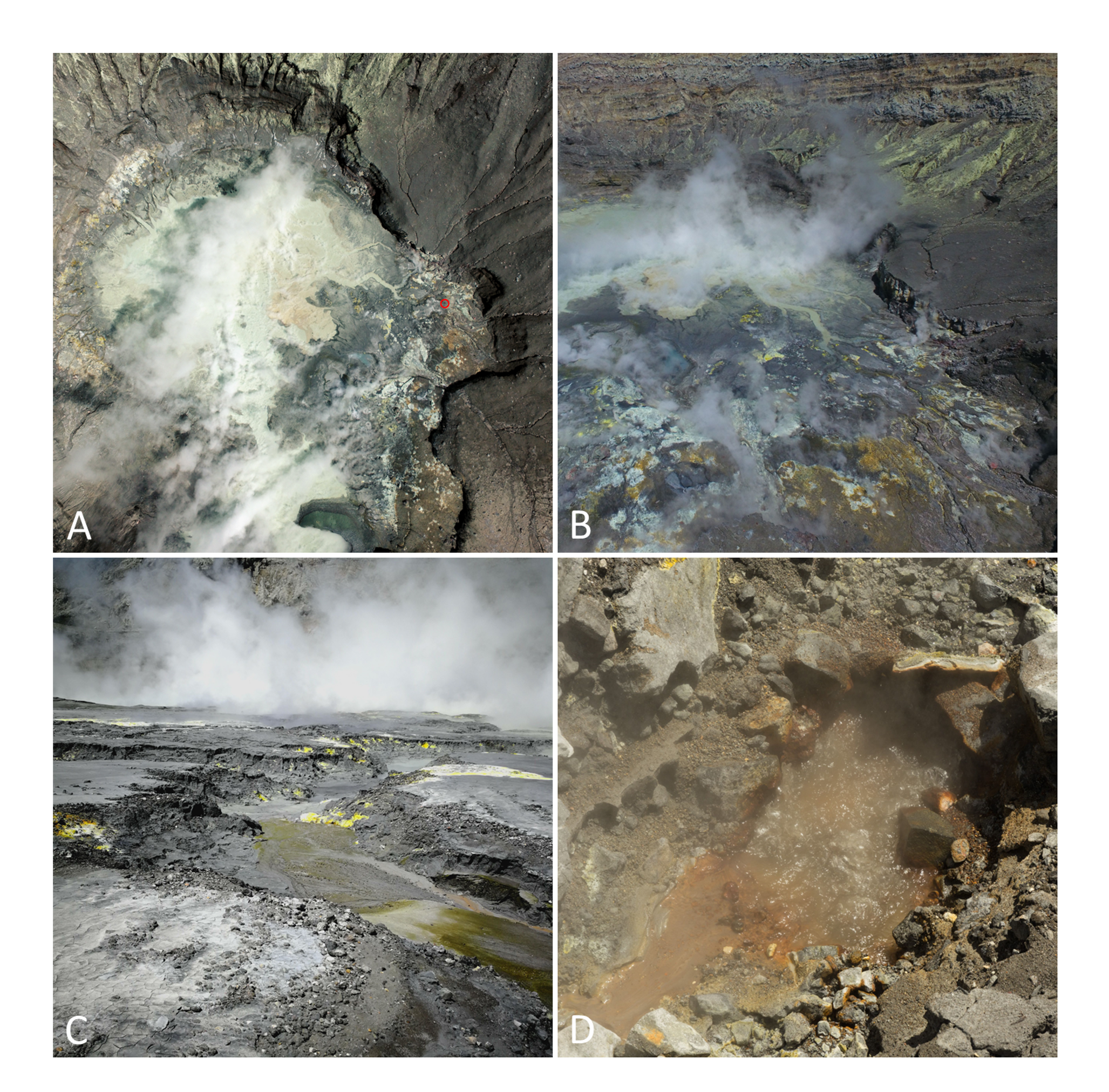

### Supp. Figure S2

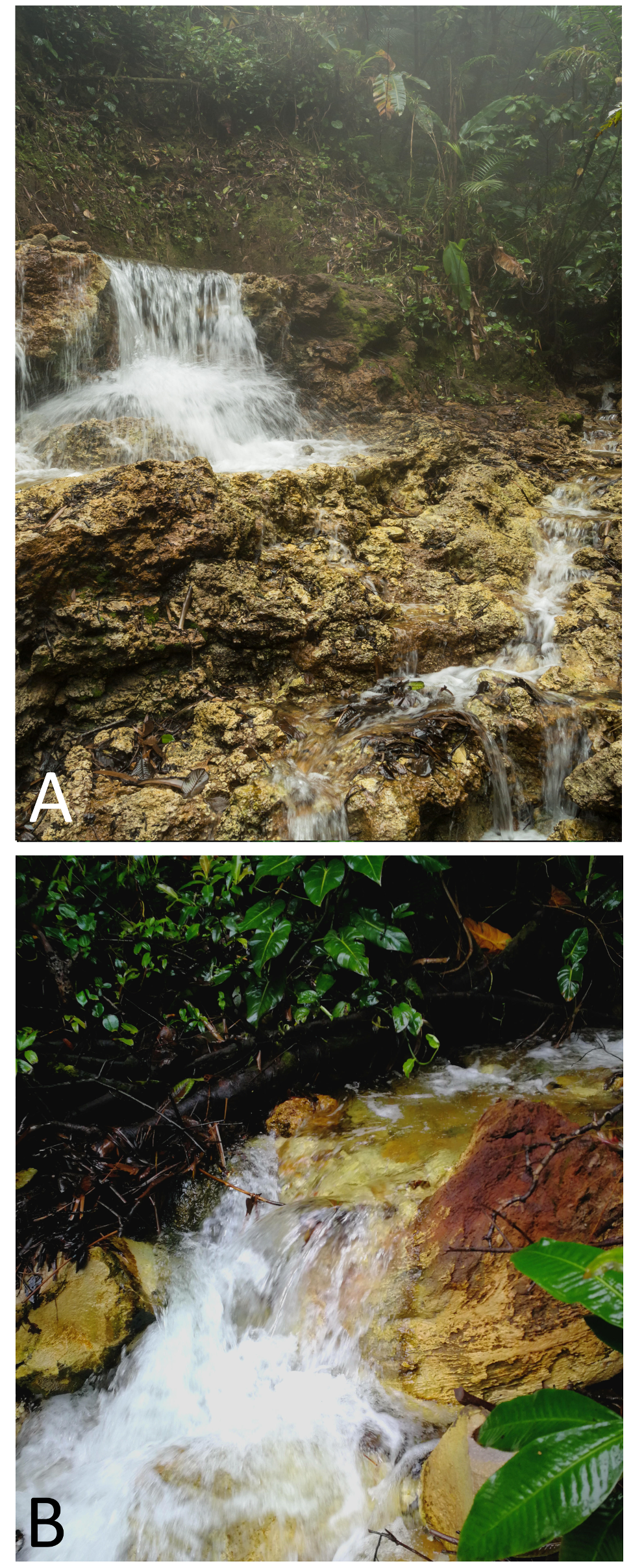

### Supp. Figure S3

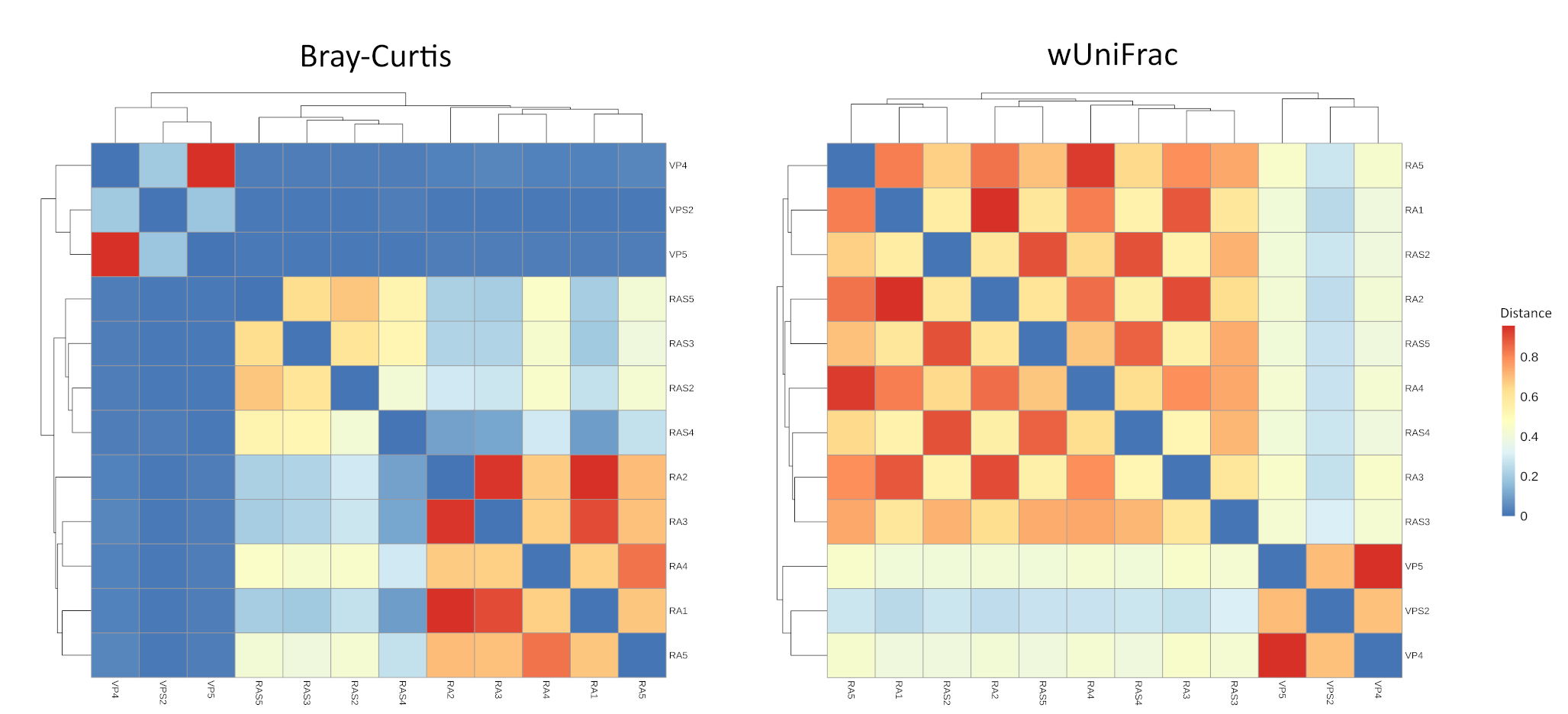

### Supp. Figure S4

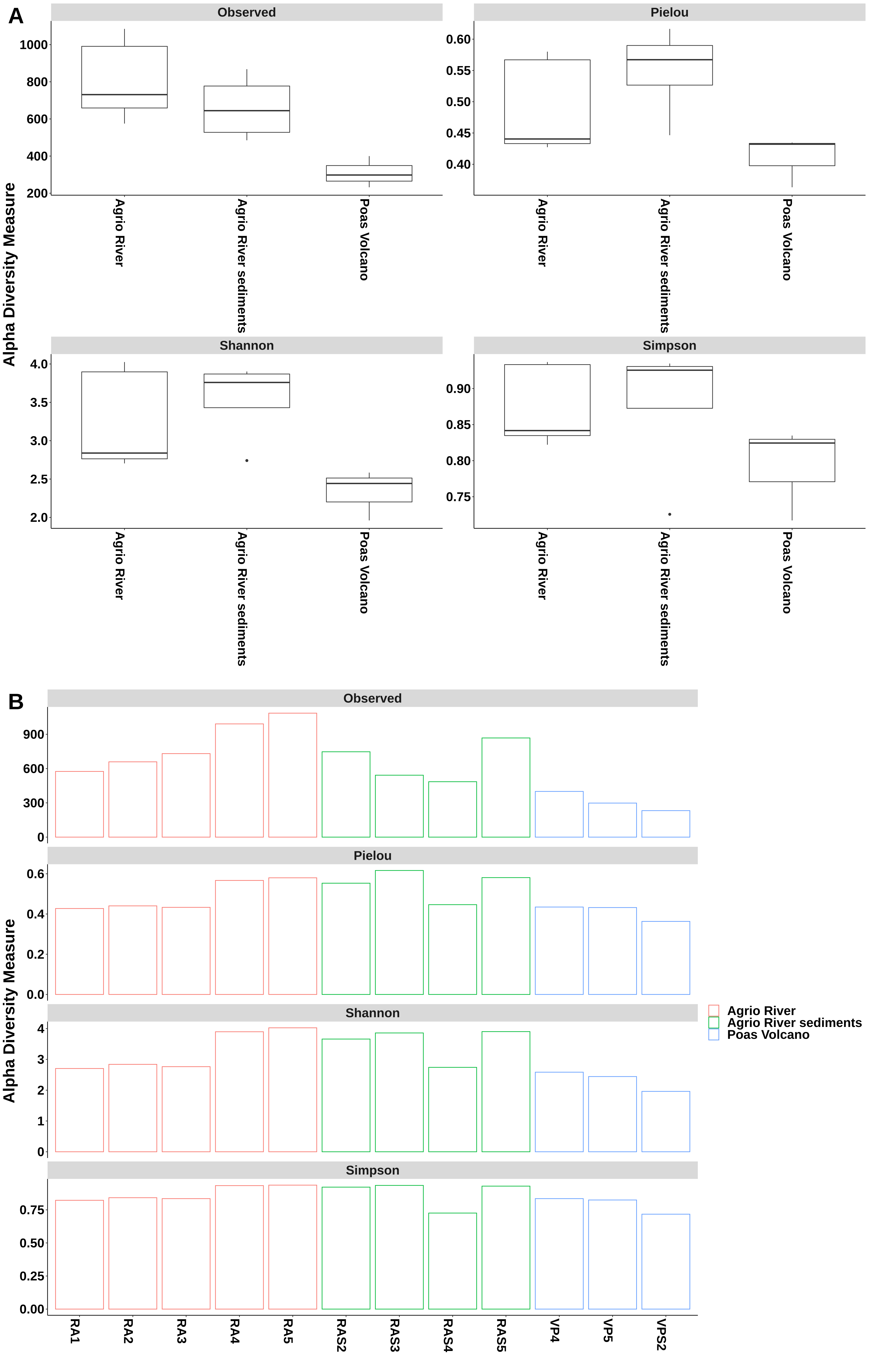
